# mtDNA diversity in human populations highlights the merit of haplotype matching in gene therapies

**DOI:** 10.1101/072348

**Authors:** E.C. Røyrvik, J.P. Burgstaller, I.G. Johnston

## Abstract

Modern gene therapies aim to prevent the inheritance of mutant mitochondrial DNA (mtDNA) from mother to offspring by using a third-party mtDNA background. Technological limitations mean that these therapies may result in a small amount of maternal mtDNA admixed with a majority of third-party mtDNA. This situation is unstable if the mother’s mtDNA experiences a proliferative advantage over the third-party mtDNA, in which case the efficacy of the therapy may be undermined. Animal models suggest that the likelihood of such a proliferative advantage increases with increasing genetic distance between mother and third-party mtDNA, but in real therapeutic contexts the genetic distance, and so the importance of this effect, remains unclear. Here we harness a large volume of available human mtDNA data to model random sampling of mother and third-party mtDNAs from real human populations. We show that even within the same haplogroup, genetic differences around 20-80 SNPs are common between mtDNAs. These values are sufficient to lead to substantial segregation in murine models, over an organismal lifetime, even given low starting heteroplasmy, inducing increases from 5% to 35% over one year. Randomly pairing mothers and third-party women in clinical contexts thus runs the risk that substantial mtDNA segregation will compromise the beneficial effects of the therapy. We suggest that choices of ‘mtDNA donors’ be based on recent shared maternal ancestry, or, preferentially, explicit haplotype matching, in order to reduce the potential for problems in the implementation of these therapies.

## Introduction

Mitochondria are small organelles within eukaryotic cells that are vital for the normal aerobic production of ATP, the ‘universal’ biochemical energy carrier. Each mitochondrion, of which there are many in any given cell, carries at least one copy of its own, small genome (mitochondrial or mtDNA), distinct from the large genome stored in the nucleus. While there are good reasons for retaining some genes in the mitochondrion (Johnston and Williams, 2016), a challenging biochemical environment and comparative lack of efficient DNA repair mechanisms allows a higher mutation rate there than in the nucleus (Alexeyev *et al.*, 2013).

Differences in the sequence of mitochondrial DNA can arise at the level of individuals (population diversity) or different mitochondria in the same cell (*heteroplasmy* – see below). In humans, mtDNA is inherited uniparentally, via the mother’s egg cell; recombination is usually negligible between human mtDNAs (Hagelberg, 2003, Hagstrom *et al.*, 2014). Given the non-recombining nature of the mitochondrial genome, such polymorphisms as exist can be expressed in terms of a straightforward phylogenetic tree (see Fig. 1A). The sum of polymorphisms in an mtDNA sequence is known as a haplotype, and any hierarchical clade of haplotypes is a haplogroup. Since inheritance is uniparental, mtDNA haplogroups are strongly susceptible to genetic drift, and this has given rise to pronounced haplogroup pattern differences between geographical areas, especially on a continental scale (see Fig. 1B).

**Figure 1.**
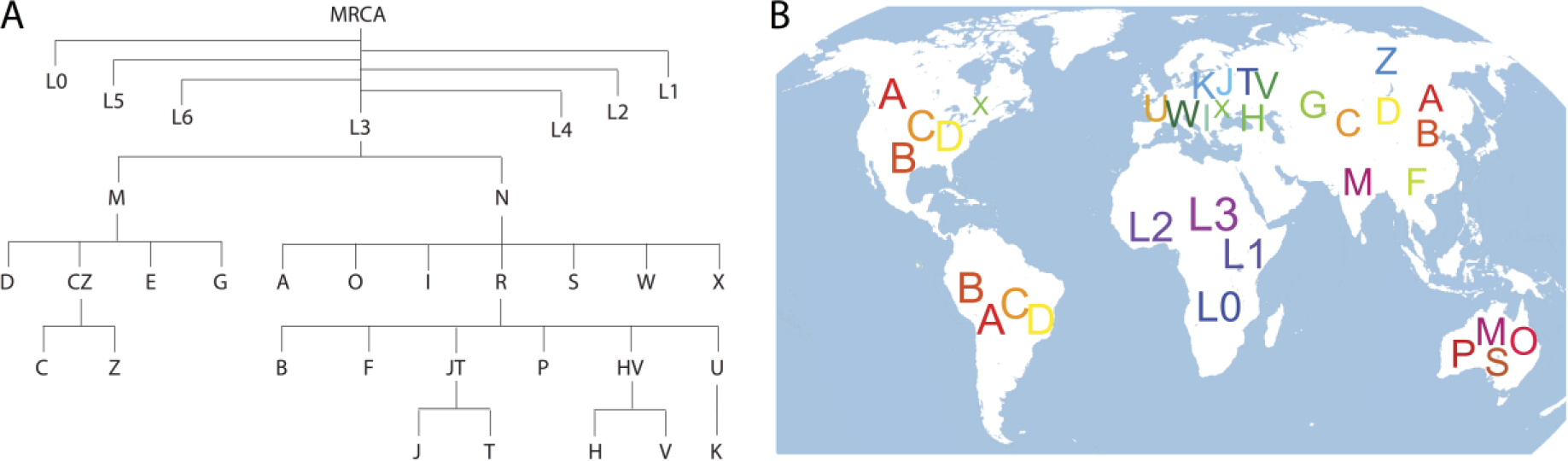
**A) Relationship between human mtDNA haplogroups.** Haplogroup labels and tree structure for human mtDNA groups; *MRCA* is most recent common ancestor. **B) Typical haplogroups in pre-colonial human populations by approximate geography.**We have omitted higher-order haplogroups of which many sub-groups are presented (e.g. *N* & *R*). Based on data from MitoMAP (Lott *et al.*, 2013) and references therein.

While most mitochondrial diversity in humans is neutral or near-neutral (Chinnery and Hudson, 2013), certain mtDNA mutations in humans can cause fatal, incurable diseases (for example, mt3243A>G, causing the inherited disease MELAS), often manifesting when the proportion of mutated mtDNA molecules in a cellular population exceeds a threshold (Taylor and Turnbull, 2005, Wallace and Chalkia, 2013). Clinical approaches to prevent the inheritance of diseases resulting from damaging mutations in mtDNA are a focus of current medical research. Cutting-edge therapies including pronuclear transfer and chromosomal spindle transfer attempt to address the inheritance of mutant mtDNA from a maternal carrier by transferring the nuclear genome (either as the pair of pronuclei or the chromosomal spindle) into an enucleated third-party oocyte or enucleated zygote with non-pathogenic mtDNA (Brown *et al.*, 2006, Burgstaller *et al.*, 2015, Craven *et al.*, 2010, Tachibana *et al.*, 2009) (Fig. 2). These therapies thus aim to place parental nuclear DNA on a healthy mitochondrial background with no mtDNA from the mother present. However, technological limitations currently mean that *carryover* is possible, whereby some of the mother’s mtDNA may be carried into the third-party cell with the transferred nuclear genetic material. These therapies can thus lead to the coexistence of several distinct sequences within cellular mtDNA populations. First, the non-pathogenic mtDNA from the third-party oocyte donor is present. Second, due to carryover, non-pathogenic mtDNA from the mother may be present. Third, due to carryover, pathogenic (mutant) mtDNA from the mother may be present (Fig. 2).

**Figure 2.**
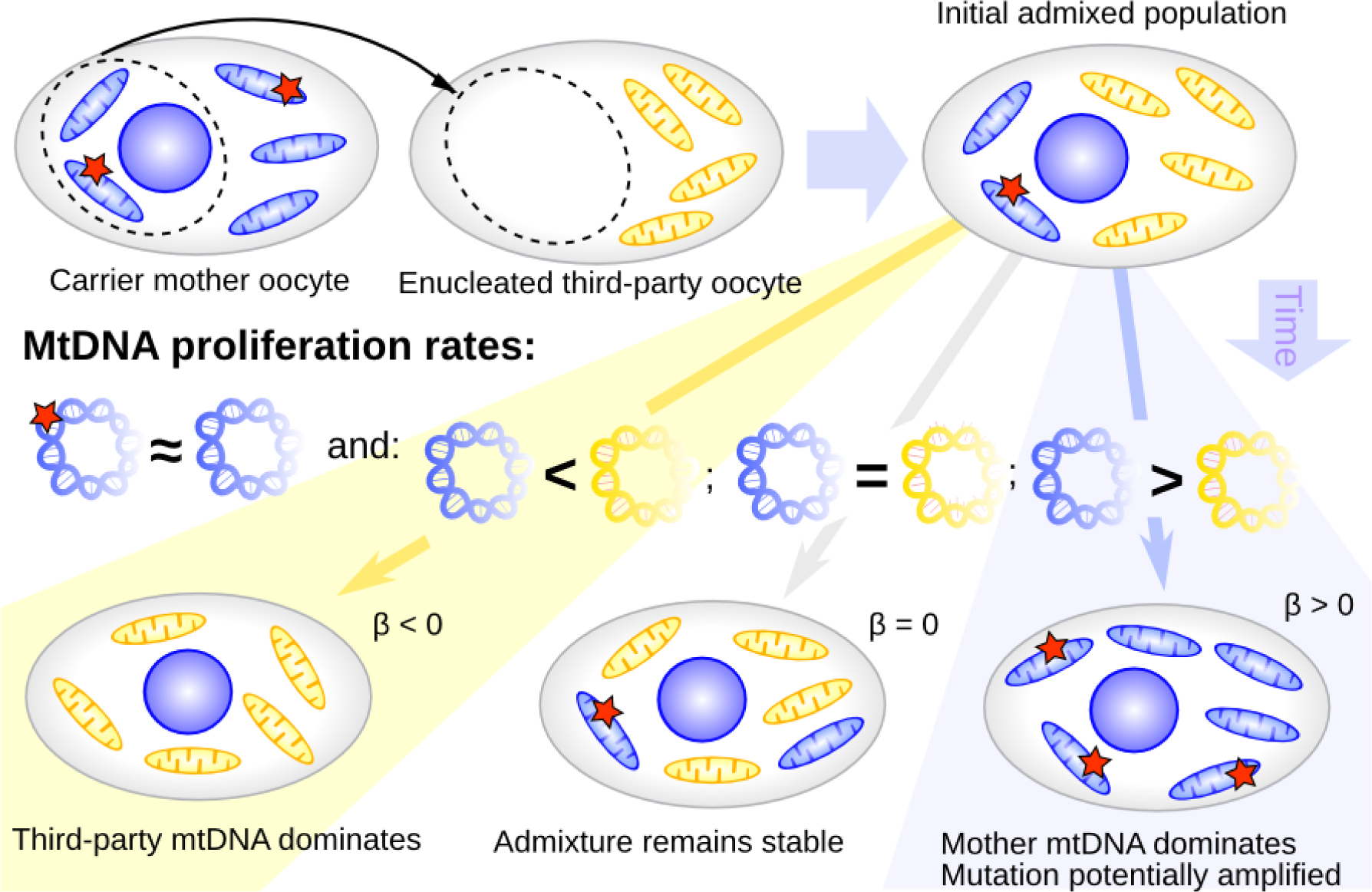
mtDNA segregation and gene therapies. A mother may possess two similar haplotypes, one wild type (blue) and one mutant (blue with red star). Therapies attempt to use a third-party with a potentially different mtDNA haplotype (yellow) to provide a healthy mtDNA background. Carryover in these therapies may result in an admixture of wildtype mother, mutant mother, and wildtype third-party mtDNA in a cell. If the two haplotypes (blue and yellow) proliferate differently, the offspring may evolve a predominance of third-party (lower left) or mother (lower right) mtDNA with time. In the latter case, if mutated mtDNA proliferates at a similar rate to its ‘carrier’ haplotype, the damaging mutation may be amplified to harmful levels in cells.

This admixture is stable if mother and oocyte donor mtDNA experience no proliferative differences (Fig. 2, centre), and if the oocyte donor haplotype experiences a proliferative advantage then carried-over mtDNA will generally be reduced over time (Fig. 2 left). However, a general proliferative advantage of the mother’s haplotype can in principle lead to the amplification of the associated pathological mutation, working against the desired effect of the therapy to remove this mutation (Fig. 2 right). This amplification can in principle occur even if the pathological mutation experiences a selective disadvantage – if this disadvantage is of lower magnitude than the proliferative difference between haplotypes, the latter effect will still dominate.

It has been shown that in cellular admixtures in mice (and other species), such proliferative differences between haplotypes do indeed commonly exist (Fig. 2; a selection of models and studies exhibiting this behaviour is given in (Burgstaller *et al.*, 2015). It has also been demonstrated that, while the direction and tissue-dependence of this differential proliferation are currently difficult to predict, its expected magnitude depends on the genetic difference between haplotypes (Burgstaller *et al.*, 2014) (Fig. 3). An important question to consider in gene therapies is thus, given the mtDNA diversity in human populations, what genetic differences are likely to arise in nuclear mother-oocyte donor pairings in therapeutic contexts, and are the proliferative differences in Fig. 2 thus likely to arise?

**Figure 3.**
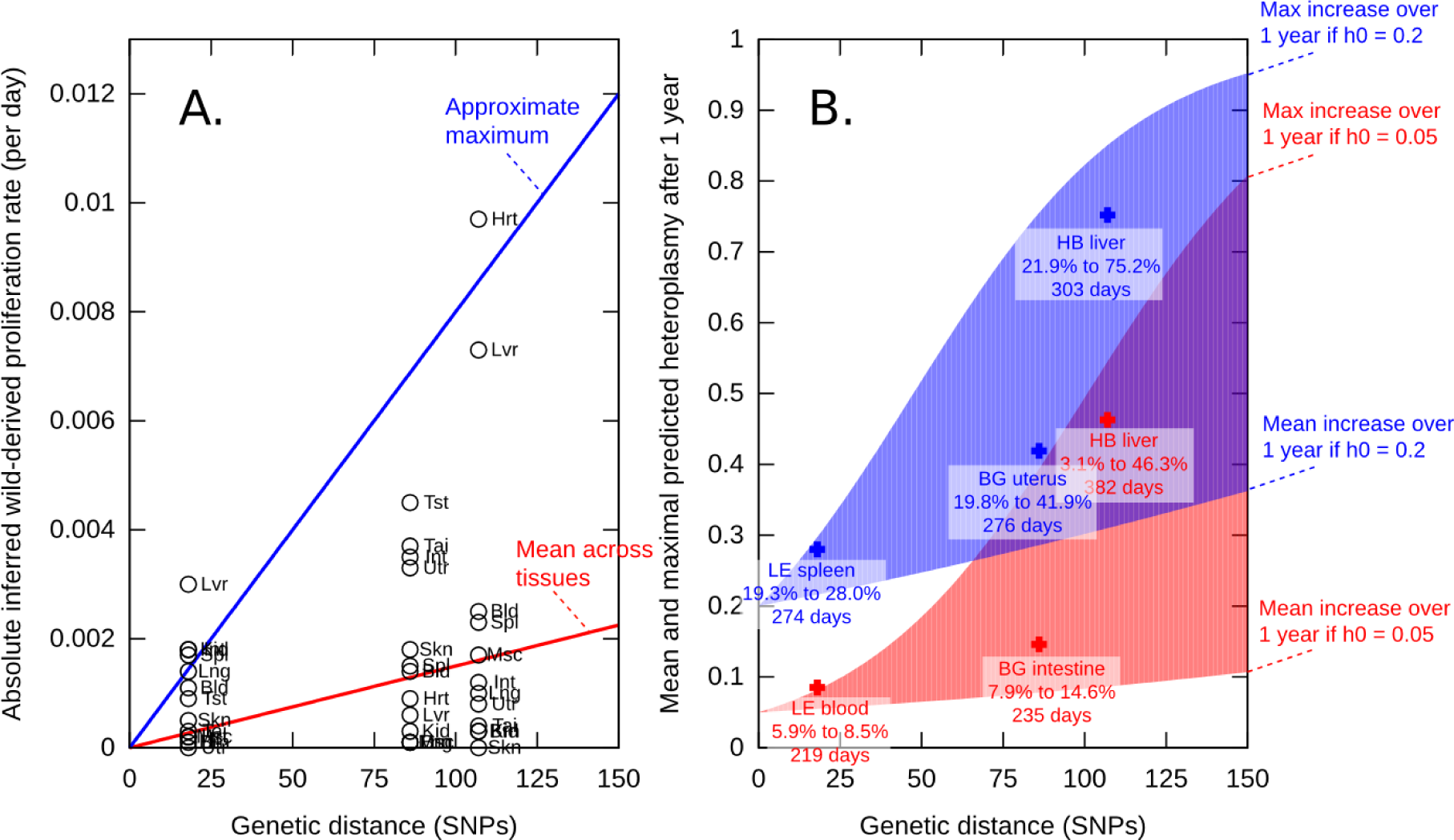
mtDNA segregation and genetic differences in mice. **A)** Magnitudes of segregation (proliferative differences between mtDNA types) in different tissues (points) in four different mtDNA pairings from (Burgstaller *et al.*, 2014). More pronounced segregation is observed in those pairings with the greatest genetic distance. Red line shows the mean trend of segregation with number of nucleotide differences; blue line shows the approximate maximum segregation strength across all tissues for mtDNA pairings with < 100 nucleotide differences. **B)** Ranges of expected heteroplasmy in mice after 1 year, given different initial heteroplasmies (h_0_) and the mean (lower) and maximal (higher) segregation magnitude observed in mice. For example, the darker red curve shows that for an mtDNA pairing with 75 nucleotide differences, a maximal increase from *h* = 0.05 to ≃ 0.3 is expected.

If Π_*ij*_ is the number of non-identical bases between two mtDNA genomes, *i* and *j*, then, intuitively, identical mtDNAs (Π_ij_ = 0) would be expected to behave identically, but the more different the mtDNAs (Π_ij_ > 0), the larger is the proliferative difference generally expected between the two. We define *heteroplasmy*, *h*, as the proportion of one ‘foreign’ mtDNA haplotype in a cellular admixture: hence, if a cell contains *H*_0_ mtDNAs of its ‘native’ haplotype and *H*_1_ mtDNAs of a ‘foreign’ haplotype, *h* = *H*_1_/(*H*_0_ + *H*_1_).

Proliferative differences between haplotypes can be measured as a quantity *β* a rate of proliferation of one mtDNA over another. In this picture, heteroplasmy *h(t)* is a function of an initial heteroplasmy *h(t=0)* and time *t*, with β describing the rate of change of heteroplasmy: 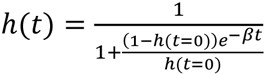. High *β* values correspond to high proliferative differences (*H*_1_ dominating over *H*_0_), and hence high probabilities of amplification of one mtDNA type with time. For example, proliferative differences of average magnitude |*β*| ≃ 0.008 per day have been measured between two mtDNA types of Π_ij_ ≃ 100 in the livers of mice; this value of *β* corresponds to an amplification of *h* from 0.05 (5% of one haplotype) to 0.49 (49% of that same haplotype) over one year (Burgstaller *et al.*, 2014).

A subset of recent evidence for proliferative differences between mtDNA haplotypes in mice is shown in Fig. 3. Fig. 3A shows inferred values of |*β*|, and the magnitude of proliferative differences between mtDNAs, in a variety of tissues for three mtDNA pairs (where Π_*ij*_ = 18, 86, and 107). Fig. 3B shows the predictions that this behaviour of *β* makes about absolute changes in heteroplasmy, for two putative admixtures beginning with 5% and 20% of a ‘foreign’ haplotype. For example, a haplotype differing from the ‘native’ type by Π_*ij*_ ≃ 100 may readily experience amplification from 5% to 50% over one year.

For simplicity, these plots are limited to the behaviour over one year, but the trends are observed to continue throughout organismal lifetimes. For example, one observation in (Burgstaller *et al.*, 2014) showed heteroplasmy in liver tissue rising from 5.9% to 81.8% over 680 days for a particular mtDNA pairing where Π_*ij*_ = 108. There is thus evidence that, in mice, nucleotide differences around Π_*ij*_ ∼ 100 are associated with proliferative differences capable of amplifying an admixed haplotype from a 5% minority to a pronounced cellular majority over the course of an organismal lifetime. But what are standard values of Π_*ij*_ in actual human populations? And is this magnitude of genetic diversity expected to give rise to clinically relevant mtDNA behaviour, given that a mutant mtDNA load of 40-60% is often sufficient to cause morbidity, and it still poorly known what ‘safe’ levels may be in most cases (Wallace and Chalkia, 2013)?

Existing studies have characterised the nucleotide differences in contemporary human populations, finding typical differences of dozens of nucleotides across modern Europeans (Fu *et al.*, 2012), greater diversity in Africa than in Europe (Briggs *et al.*, 2009), and results confirming and expanding these observations across a broader geographical range (Lippold *et al.*, 2014). A modern workflow has been developed to address related evolutionary questions (Blanco *et al.*, 2011). However, to our knowledge, the interpretation of these statistics in terms of mtDNA segregation possibility and implications for disease is currently absent, as is an attempt to characterise the expected diversity in modern populations combining social (census) and biological (sequence) data.

## Materials and Methods

### Materials

None.

### Methods

We took a data-driven approach, harnessing the large numbers of human mtDNA sequence data now available through the NCBI database, as well as haplogroup data in the literature. mtDNA molecules may be categorised, via the presence or absence of diagnostic SNPs, into haplogroups, which are typically designated by an alphanumeric code and follow a moderately complex hierarchy. For example, at the coarsest level, all human mtDNAs so far recorded fall into haplogroup *L*. Subsets of *L* include *N* (which in turn includes *R*, containing *H* and *V, etc.*) and *W*, *X*, *Y* and others. A simplified tree of haplogroups is shown in Fig. 1A and illustrative geographical distributions are shown in Fig. 1B.

Data on the haplogroup makeup of ‘pre-colonial populations’, i.e. before early modern population mixing, from different geographical regions is available via MitoMAP (Lott *et al.*, 2013). These data can be used to estimate the probability that an individual with maternal ancestry from a given region belongs to a given haplogroup.

Many specific mtDNA sequences corresponding to individual humans belonging to a given haplogroup are available via NCBI. Using these data, we sought to identify the expected genetic differences between pairs of individual, real human mtDNAs. To estimate these expected differences, we first characterised the expected differences between specific mtDNA samples within and between different haplogroups. For the purposes of this study we employ the convention that an individual is marked as a member of haplogroup category ℋ if it is (a) a member of ℋ and (b) **not** a member of any haplogroup that is a subgroup of ℋ. For example, *x* is labelled as *L* if *x* is in haplogroup *L* but not in any of the *A, B, C, D, E*,… that are subgroups of *L*.

We obtained the > 30*k* mtDNA sequences available from NCBI Nucleotide database (NCBI, 2015). Of these sequences ∼ 7.6*k* had straightforwardly interpretable haplogroup information, where the initial letter of the */haplogroup* field was taken to be the haplogroup label. We categorised these records by this initial letter, then employed the following sampling protocol. Given a pair of haplogroups {ℋ_1_, ℋ_2_}, we picked at random a sequence belonging to ℋ_1_ and picked at random a sequence belonging to ℋ_2_ (ensuring that the two sequences were not the same sample if ℋ_1_ = ℋ_2_,). We used BLAST to record the number of sequence differences between these specific sampled sequences. For the purposes of this report we recorded the number of non-identical bases as the nucleotide difference Π_*ij*_; we also note that indels commonly exist between sampled mtDNA sequences, further contributing to mtDNA diversity. We then built up a distribution of sequence differences over many (*n=1000*) sampled pairs of specific human mtDNAs from the given pair of haplogroups.

To connect more explicitly with medical policy, we next changed the scale of our analysis from haplogroups *per se* to the estimated haplogroup profiles of real human populations. First, we employed heuristic data from the MitoMAP project (Lott *et al.*, 2013) estimating the haplogroup makeup of pre-colonial populations from different regions of the world, while noting that the actual census populations will usually have a very different makeup, especially in New World countries that experienced extensive overseas colonization. For each region, we randomly chose two haplogroups, each with a probability corresponding to that haplogroup’s representation in the region of interest. We then randomly chose two specific mtDNA sequences from those two haplogroups. As above, we then used BLAST to determine the genetic difference between those specific sequences. We repeated this process many times to build up an expected distribution of the genetic differences between two randomly chosen members of the human population from that region.

As the UK is on the cusp of implementing gene therapies based on nuclear transfer, we then performed a more rigorous, population-based analysis for Britain. In order to estimate the probable levels of nucleotide diversity (Π_*ij*_) in mtDNA between two randomly selected British women, and hence the likely magnitude of proliferative differences between their mtDNA, a haplogroup profile of Britain was assembled, based on over 4,600 individuals. The majority of the UK samples represent ethnic Britons. To account for the fact that the modern UK population consists of many ethnicities, approximations of mtDNA haplogroup distributions for the two largest cities in the UK (London and Birmingham) were also constructed. These distributions are estimates, based on data from the 2011 census, immigration data, and published mtDNA haplogroup data for areas from which there has been mass immigration into the UK (see SI for details).

For each ethnic census category, an estimate of probable haplogroup composition was created (see SI for details on calculations), and the frequency values scaled by the numerical census data to yield expected haplogroup frequencies in London and Birmingham. For simplicity, the single letter level of nomenclature is used, with the exception of superhaplogroup *L*, for which its subgroups *L0-3* are included.

## Results

Fig. 4A shows the resulting statistics on differences between sampled mtDNA sequences between haplogroup pairs. Several intuitive features are immediately observable. First, haplogroup *L* displays noticeably more intra-haplogroup differences than any other haplogroup. *L* haplogroups constitute the majority of African haplogroups (and have very deep branching times relative to non-African haplogroups) and are thus expected to include the most genetic diversity (Behar *et al.*, 2008). Second, with the exception of *L*, diagonal elements (i.e. samples from a haplogroup compared to samples from the same haplogroup) show less diversity than off-diagonal elements (i.e. samples from a haplogroup compared to samples from a different haplogroup). Third, haplogroup pairings which are expected to be similar (for example, sister clades *H* and *V*) show decreased genetic diversity. The inset shows a breakdown of the *L* haplogroup into its immediate subgroups.

**Figure 4.**
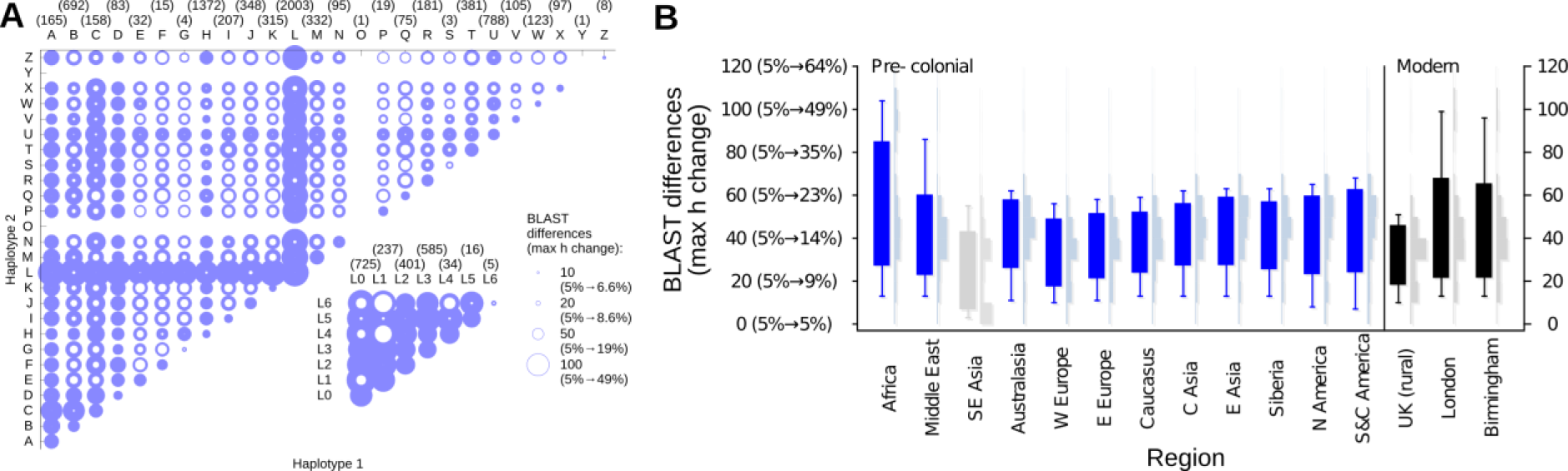
**A) MtDNA differences between haplogroups.** The maximum (outer halo) and minimum (inner halo) nucleotide differences expected between a pair of randomly sampled mtDNA sequences (horizontal and vertical axes). The diagonal corresponds to pairs within the same haplogroup; off-diagonal elements correspond to pairs of mtDNAs from different haplogroups. Dataset size for each haplogroup is given in brackets; n=1000 samples were used for each pairing. Max *h* change shows, for a given magnitude of genetic diversity, the maximum expected change in heteroplasmy over one year starting at 5%, based on mouse models (Fig 3). As described in the text, haplotype labels denote sequences that fall within a given category and not within any named subcategories of that category. Inset shows subgroups of the most-diverse *L* haplogroup. **B) MtDNA differences between geographical regions.** In blue, genetic differences between a pair of individuals randomly sampled from sets modelling populations within a given region of the world, using the MitoMAP (Lott *et al.*, 2013) estimation of the (pre-colonial) haplogroup profile of different geographical regions. In black, expected differences in the general the modern non-urban UK population, and populations of London and Birmingham. Candlesticks show minimum, mean ± s.d., and maximum nucleotide differences between simulated pairs sampled from geographical regions. Explicit sample distributions are given in in lighter colours; max h change gives maximum expected change in heteroplasmy as in (A). SE Asia (in grey) has poorly characterised MitoMAP estimates.

A notable result from this analysis is that between haplogroups, differences of ∼50 SNPs are common, and, even within haplogroups, differences of ∼20 SNPs are common. This level of diversity may not seem substantial when compared to the ∼16 kilobases of total human mtDNA, but we draw attention to our previous observations that differences of ∼ 20 SNPs were enough to induce significant proliferative differences between haplotypes in mice, who also have a ∼16kb mtDNA genome (Burgstaller *et al.*, 2014). As shown in parentheses in Fig. 4A, the magnitudes of Π that likely emerge from pairwise haplotype samples match those responsible for dramatic mtDNA heteroplasmy changes in mouse models.

Fig. 4A also provides a means of identifying a ‘partner’ for a given haplogroup that minimizes Π and hence the likelihood of damaging segregation. For example, given a mother with haplogroup B and a choice between donors from C, V, and L, Fig.4A shows that the B-V pairing minimizes maximum Π, and thus affords the lowest risk of high segregation (see Discussion).

Table 1 gives the estimated haplogroup makeup of the UK and two major cities, based on a combination of census and immigration data and a survey of worldwide mtDNA sequences (see Methods and SI). We underline that these quantities are principled estimates, but the summary statistics that arise from these estimates are robust to variation in the exact population frequencies, and is consistent with the behaviour expected from an ethnically mixed population based on more direct estimates (see below).

**Table 1.** Estimated haplogroup frequencies in the British population. UK – majority ethnic Britons, exclusive of large urban areas, London, Birmingham –census and immigration data based estimates (see SI).

| HG | UK % | London % | Birmingham % |
| --- | --- | --- | --- |
| A | 0.0% | 0.6% | 0.5% |
| B | 0.0% | 1.1% | 0.7% |
| C | 0.0% | 0.3% | 0.2% |
| D | 0.0% | 0.8% | 0.5% |
| F | 0.0% | 1.1% | 0.8% |
| G | 0.0% | 0.2% | 0.2% |
| H | 45.2% | 30.4% | 29.9% |
| I | 4.1% | 2.6% | 2.6% |
| J | 12.4% | 7.8% | 8.2% |
| K | 8.3% | 5.1% | 5.3% |
| L0 | 0.0% | 1.3% | 0.8% |
| L1 | 0.0% | 2.4% | 1.9% |
| L2 | 0.0% | 4.9% | 3.8% |
| L3 | 0.1% | 4.5% | 3.5% |
| M | 0.0% | 10.4% | 12.7% |
| N | 0.0% | 0.1% | 0.2% |
| O | 0.0% | 0.0% | 0.0% |
| P | 0.0% | 0.0% | 0.0% |
| R | 0.1% | 2.7% | 3.6% |
| S | 0.0% | 0.0% | 0.0% |
| T | 10.5% | 6.8% | 6.9% |
| U | 12.6% | 11.5% | 12.6% |
| V | 3.2% | 1.6% | 1.8% |
| W | 1.5% | 1.2% | 1.6% |
| X | 1.8% | 1.2% | 1.1% |
| other | 0.3% | 1.3% | 0.6% |

Fig. 4B illustrates the distribution of nucleotide differences between individuals sampled from geographical regions, and rural vs. urban UK based on estimates in Table 1, in this manner. It is immediately noticeable that pairs of individuals from Africa generally exhibit more diversity than pairs chosen from other regions, but it is striking that the expected genetic difference in many geographic regions is around Π_*ij*_ ∼ 40-50 SNPs, often with a range between 10-100 SNPs. The substantial diversity expected in the UK and its cities is of a consistent magnitude with that expected from its population history, involving admixtures of African and Asian immigrants in addition to its original European state. Again, parenthesized numbers in Fig. 4B illustrate that these magnitudes of Π are readily able to induce pronounced heteroplasmy shifts in mice. Taken together, these results demonstrate that expected levels of mtDNA diversity in modern human populations are of comparable magnitude to those responsible for substantial segregation bias in a existing mammalian models, and so therapies that randomly pair women from these populations may be subject to potentially detrimental heteroplasmy changes over time.

## Discussion

Our analysis clearly shows that, even within a geographical region restricted to the point of being dominated by a single mtDNA haplogroup, a Π_*ij*_ = 10 − 100 is expected between randomly sampled individuals from that region. On a continental scale, expected differences are highest in Africa, as predicted from our knowledge of human population history, and comparably lower elsewhere. Comparably high, however, are the differences in the largest urban populations of the UK, where oocyte donor therapies will be implemented.

In mice, proliferative differences between haplogroups with Π_*ij*_ ∼ 100 were sufficient in some tissues to cause amplification of one mtDNA type from 0.05 to 0.64 (i.e. a small representation to a notable majority) over an organismal lifetime (Fig. 2B). There remains a wide range of questions involving the mapping from the murine model to the human system. One criticism of our argument may be that mtDNA segregation in humans may progress more slowly than in mice, reducing the magnitude of the effects we consider. However, segregation in humans has been observed to occur more rapidly than in mice (Wallace and Chalkia, 2013). Furthermore, evidence exists for pronounced segregation of a pathological mutation over very short times during embryo-fetal development (Monnot *et al.*, 2011), suggesting the presence of mechanisms in humans that support fast segregation, and which could in principle also act on non-pathological mutations. Even in a conservative picture where mtDNA turnover rates are scaled by organismal lifetimes, amplification over the (longer) human lifetime will still be anticipated by analogy with the murine system. An important clinical example of the potentially high mtDNA segregation in human disease (again involving a pathological mutation) is described in Ref.(Mitalipov *et al.*, 2014), in which an embryo selected for its low (12%) load of the 3243 mutation (Treff *et al.*, 2012) developed into an infant with >40% loads in blood and urine at six weeks of age, presenting with a range of (possibly unrelated) metabolic pathologies.

It is worth noting that, in addition to the unpredictability of segregation direction, the rate at which mtDNA segregation occurs is not simple and constant – rather, it can depend on tissue type, organismal age and developmental stage (Burgstaller *et al.*, 2014), and complicating processes including the mtDNA bottleneck (Johnston *et al.*, 2015). It is thus not unreasonable to think that the ‘averaged’ rates reported here may be underestimates for a particular time period. We therefore highlight that, even from a conservative calculation of segregation rates, *the likely genetic differences between humans randomly sampled from a population may well allow substantial amplification of a disease-carrying mtDNA haplotype over the timescale of a human lifetime*.

We must also consider whether randomly sampling NCBI sequences is a good model for the mtDNA pairings likely to be involved in gene therapies. The counterexample of this would be a population consisting of many individuals with identical mtDNA sequences and a small number of individuals with different sequences. The NCBI, which assigns records to unique sequences, will likely have one record for the common sequence and one each for the rare different sequences. In this case, uniformly sampling the NCBI would underestimate the population fraction with the common sequence, and thus tend to overestimate mtDNA diversity. However, the ubiquity of many-SNP differences between records (see Fig.4) suggests that this problematic population structure is unlikely, and indeed, several contemporary studies have observed differences between each individual sample (Fu *et al.*, 2012, Lippold *et al.*, 2014). Additionally, socio-economic factors will give rise to structure in the pairings in clinical applications (which may either decrease or increase the expected Π_*ij*_). Despite these complications, we consider our approximations appropriate for considering first-order bounds of likely behaviour in these populations exhibiting realistic human diversity.

The danger of pathological mutations ‘hitchhiking’ on favoured haplotypes and being amplified along with the haplotype is described in the introduction and has been discussed previously (Burgstaller *et al.*, 2014, Burgstaller *et al.*, 2015). An additional danger is the amplification of an initially rare mtDNA haplotype to the point where it competes with the dominant mtDNA type in a cell and causes pathologies through mismatched protein subunits or other mechanisms (Burgstaller *et al.*, 2015). The co-occurrence in a cell of two different, but both separately non-pathogenic, mtDNAs has been observed to result in adverse physiological changes (Sharpley *et al.*, 2012). Segregation between mtDNA haplotypes, allowing an initially rare haplotype to proliferate and become amplified within a cell, has the potential to manifest and exacerbate both of these potential issues.

To diminish the likelihood of potentially harmful mtDNA segregation, which we argue is likely given the mtDNA diversity in the modern UK population, we urge experts involved in the implementation of these therapies to consider ‘haplotype matching’, i.e. choosing an oocyte donor with mtDNA as similar as possible to the mother’s in clinical approaches. Methods to match haplotypes (minimise Π_*ij*_) could include choosing maternal relatives of the mother with low or zero proportions of the pathological mutation under consideration, or choosing donors from a haplogroup as similar as possible to the mother’s. To illustrate this latter strategy, Fig. 5 shows the range of expected Π values that could arise when a third-party donor is paired with a mother from haplogroup H1a. If no haplotype matching is employed, and the third-party donor is randomly sampled from our estimated London population, a maximum Π around 100 is possible (due to the pronounced population diversity illustrated in Fig. 4B). Choosing a donor from haplogroup H decreases this maximum value to around 36 (that is, the maximal within-H diversity, shown on the diagonal of Fig. 4A). More detailed matching, specifically choosing another H1a woman as the third-party, further limits the maximum Π to approximately 17. These lower values achieved through haplotype matching dramatically decrease the expected potential heteroplasmy changes (for example, in mice (Fig 2), from a maximum of 5% → 49% over one year for Π = 100 to 5% → 8% over one year for Π = 17), thus immediately limiting the potential for detrimental segregation. Our results, and future findings from more detailed studies, can help provide a strategy for this matching process – given a mother of known mtDNA haplogroup, choose from available oocyte donors so as to minimise the maximum genetic distance given in Fig. 4. Such haplotype matching, which is in principle technically straightforward and economically marginal, decreases the risk of inadvertently choosing an mtDNA pairing which experiences substantial proliferative differences, and thus decreases the risk of manifestation of the disease the therapy was implemented to prevent.

**Figure 5.**
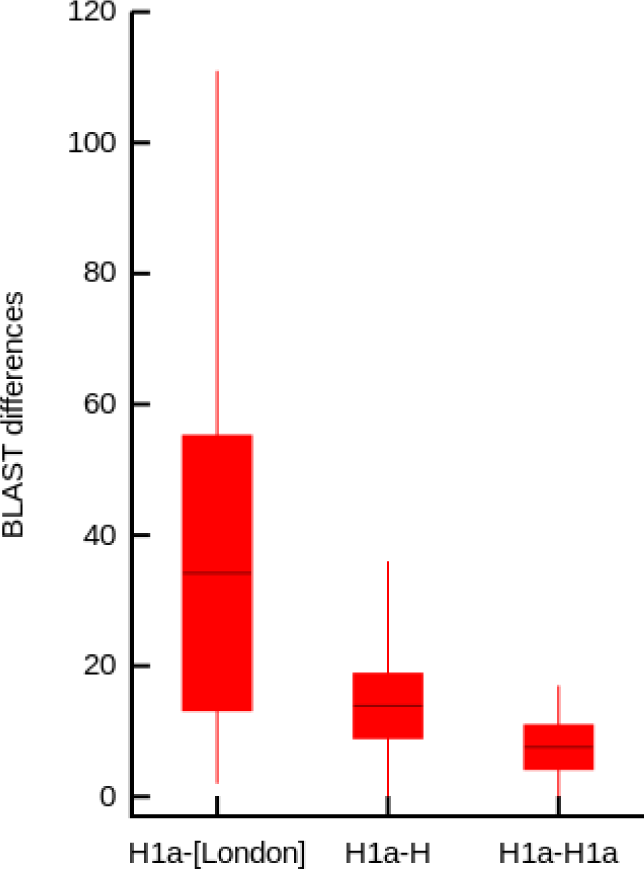
MtDNA differences expected with different haplotype matching strategies for a mother with haplogroup H1a. >Distributions of nucleotide differences (min, mean +− sd, max) expected when pairing mtDNA from haplogroup H1a with randomly sampled mtDNA from our estimated London population, with randomly sampled mtDNA from haplogroup H, and with randomly sampled mtDNA from haplogroup H1a.

## Supplementary Information

### I. UK city mtDNA haplogroup compositions

In order to produce a realistic approximation of the mitochondrial haplogroup composition of the two largest UK cities, London and Birmingham, an approach based on ethnic categories in the 2011 UK census and mt haplogroup data from corresponding regions and/or countries was employed.

Ethnic composition data were downloaded from http://webarchive.nationalarchives.gov.uk, in which 18 categories were defined (see Table 1 for these category definitions and their frequencies in London and Birmingham).

**Table 1.** Proportion of different census categories in London and Birmingham.

| Code | Category | Sub-category | London (%) | B'ham (%) |
| --- | --- | --- | --- | --- |
| Pop1 | White | English/Welsh/Scottish/Northern Irish/British | 44.8 | 53.1 |
| Pop2 | White | Irish | 2.2 | 2.1 |
| Pop3 | White | Gypsy or Irish Traveller | 0.1 | 0 |
| Pop4 | White | Other White | 12.6 | 2.7 |
| Pop5 | Mixed/multiple ethnic group | White and Black Caribbean | 1.5 | 2.3 |
| Pop6 | Mixed/multiple ethnic group | White and Black African | 0.8 | 0.3 |
| Pop7 | Mixed/multiple ethnic group | White and Asian | 1.2 | 1 |
| Pop8 | Mixed/multiple ethnic group | Other Mixed | 1.4 | 0.8 |
| Pop9 | Asian/Asian British | Indian | 6.6 | 6 |
| Pop10 | Asian/Asian British | Pakistani | 2.7 | 13.5 |
| Pop11 | Asian/Asian British | Bangladeshi | 2.7 | 3 |
| Pop12 | Asian/Asian British | Chinese | 1.5 | 1.2 |
| Pop13 | Asian/Asian British | Other Asian | 4.9 | 2.9 |
| Pop14 | Black/African/Caribbean/Black British | African | 7 | 2.8 |
| Pop15 | Black/African/Caribbean/Black British | Caribbean | 4.2 | 4.4 |
| Pop16 | Black/African/Caribbean/Black British | Other | 2.1 | 1 |
| Pop17 | Other ethnic group | Arab | 1.3 | 1 |
| Pop18 | Other ethnic group | Any other ethnic group | 2.1 | 1 |

Then, for each ethnic category suitable mtDNA datasets were collated (see Table 2 for resultant haplogroup profiles).

**Table 2.** Estimated haplogroup frequencies in UK census categories.

| HG | Pop 1 | Pop 2 | Pop 3 | Pop 4 | Pop 5 | Pop 6 | Pop 7 | Pop 8 | Pop 9 | Pop 10 | Pop 11 | Pop 12 | Pop 13 | Pop 14 | Pop 15 | Pop 16 | Pop 17 | Pop 18 |
| --- | --- | --- | --- | --- | --- | --- | --- | --- | --- | --- | --- | --- | --- | --- | --- | --- | --- | --- |
| A | 0.0 % | 0.0 % | 0.0 % | 0.0 % | 0.1 % | 0.0 % | 1.4 % | n/a | 0.0 % | 0.1 % | 10.0 % | 5.3 % | 2.7 % | 0.0 % | 0.3 % | 0.1 % | 0.1 % | n/a |
| B | 0.0 % | 0.0 % | 0.0 % | 0.0 % | 0.1 % | 0.0 % | 5.2 % | n/a | 0.2 % | 0.4 % | 0.0 % | 20.5 % | 10.4 % | 0.0 % | 0.3 % | 0.1 % | 0.0 % | n/a |
| C | 0.0 % | 0.0 % | 0.0 % | 0.0 % | 0.0 % | 0.0 % | 1.1 % | n/a | 0.5 % | 0.0 % | 0.0 % | 3.8 % | 2.1 % | 0.0 % | 0.0 % | 0.0 % | 0.0 % | n/a |
| D | 0.0 % | 0.0 % | 0.0 % | 0.0 % | 0.3 % | 0.0 % | 3.9 % | n/a | 0.1 % | 0.0 % | 0.0 % | 15.3 % | 7.7 % | 0.0 % | 0.5 % | 0.3 % | 0.0 % | n/a |
| F | 0.0 % | 0.0 % | 0.0 % | 0.0 % | 0.0 % | 0.0 % | 4.8 % | n/a | 1.1 % | 1.2 % | 0.0 % | 18.0 % | 9.6 % | 0.0 % | 0.0 % | 0.0 % | 0.0 % | n/a |
| G | 0.0 % | 0.0 % | 0.0 % | 0.0 % | 0.0 % | 0.0 % | 0.8 % | n/a | 0.8 % | 0.6 % | 0.0 % | 2.4 % | 1.6 % | 0.0 % | 0.0 % | 0.0 % | 0.0 % | n/a |
| H | 45.2 % | 50.7 % | 50.7 % | 48.1 % | 22.7 % | 24.4 % | 23.2 % | n/a | 2.5 % | 11.2 % | 0.0 % | 0.0 % | 1.3 % | 3.6 % | 0.3 % | 1.9 % | 24.1 % | n/a |
| I | 4.1 % | 3.0 % | 3.0 % | 2.9 % | 2.0 % | 2.4 % | 2.1 % | n/a | 0.2 % | 0.7 % | 0.0 % | 0.0 % | 0.1 % | 0.8 % | 0.0 % | 0.4 % | 1.4 % | n/a |
| J | 12.4 % | 11.0 % | 11.0 % | 9.2 % | 6.4 % | 6.5 % | 6.5 % | n/a | 1.5 % | 3.5 % | 0.0 % | 0.0 % | 0.7 % | 0.6 % | 0.5 % | 0.6 % | 11.2 % | n/a |
| K | 8.3 % | 12.3 % | 12.3 % | 5.2 % | 4.1 % | 4.4 % | 4.3 % | n/a | 0.7 % | 1.1 % | 0.0 % | 0.0 % | 0.3 % | 0.5 % | 0.0 % | 0.2 % | 5.4 % | n/a |
| L0 | 0.0 % | 0.0 % | 0.0 % | 0.0 % | 2.5 % | 5.8 % | 0.0 % | n/a | 0.0 % | 0.1 % | 0.0 % | 0.0 % | 0.0 % | 11.6 % | 5.0 % | 8.3 % | 2.1 % | n/a |
| L1 | 0.0 % | 0.0 % | 0.0 % | 0.0 % | 9.8 % | 7.2 % | 0.0 % | n/a | 0.0 % | 0.0 % | 0.0 % | 0.0 % | 0.0 % | 14.3 % | 19.5 % | 17.0 % | 1.9 % | n/a |
| L2 | 0.0 % | 0.0 % | 0.0 % | 0.0 % | 19.4 % | 15.1 % | 0.0 % | n/a | 0.0 % | 0.5 % | 0.0 % | 0.0 % | 0.0 % | 29.9 % | 38.8 % | 34.5 % | 4.7 % | n/a |
| L3 | 0.1 % | 0.0 % | 0.0 % | 0.0 % | 16.5 % | 14.1 % | 0.0 % | n/a | 0.0 % | 1.3 % | 0.0 % | 0.0 % | 0.0 % | 27.8 % | 33.0 % | 30.5 % | 7.8 % | n/a |
| M | 0.0 % | 0.0 % | 0.0 % | 0.0 % | 0.4 % | 1.8 % | 20.5 % | n/a | 54.3 % | 38.1 % | 66.7 % | 27.6 % | 40.9 % | 3.5 % | 0.8 % | 2.1 % | 4.0 % | n/a |
| N | 0.0 % | 0.0 % | 0.0 % | 0.0 % | 0.0 % | 0.1 % | 0.3 % | n/a | 0.1 % | 1.3 % | 0.0 % | 1.2 % | 0.6 % | 0.1 % | 0.0 % | 0.1 % | 4.3 % | n/a |
| O | 0.0 % | 0.0 % | 0.0 % | 0.0 % | 0.0 % | 0.0 % | 0.0 % | n/a | 0.0 % | 0.0 % | 0.0 % | 0.0 % | 0.0 % | 0.0 % | 0.0 % | 0.0 % | 0.0 % | n/a |
| P | 0.0 % | 0.0 % | 0.0 % | 0.0 % | 0.0 % | 0.0 % | 0.0 % | n/a | 0.0 % | 0.0 % | 0.0 % | 0.0 % | 0.0 % | 0.0 % | 0.0 % | 0.0 % | 0.0 % | n/a |
| R | 0.1 % | 0.3 % | 0.3 % | 0.0 % | 0.1 % | 0.2 % | 5.7 % | n/a | 18.0 % | 12.5 % | 6.7 % | 4.4 % | 11.2 % | 0.3 % | 0.0 % | 0.1 % | 10.4 % | n/a |
| S | 0.0 % | 0.0 % | 0.0 % | 0.0 % | 0.0 % | 0.0 % | 0.0 % | n/a | 0.0 % | 0.0 % | 0.0 % | 0.0 % | 0.0 % | 0.0 % | 0.0 % | 0.0 % | 0.0 % | n/a |
| T | 10.5 % | 7.0 % | 7.0 % | 9.7 % | 5.2 % | 5.7 % | 5.6 % | n/a | 1.3 % | 3.1 % | 0.0 % | 0.0 % | 0.6 % | 0.8 % | 0.0 % | 0.4 % | 7.2 % | n/a |
| U | 12.6 % | 12.3 % | 12.3 % | 17.0 % | 6.8 % | 7.5 % | 10.1 % | n/a | 15.4 % | 20.5 % | 16.7 % | 0.0 % | 7.7 % | 2.3 % | 1.0 % | 1.7 % | 10.3 % | n/a |
| V | 3.2 % | 0.0 % | 0.0 % | 0.1 % | 1.6 % | 1.7 % | 1.6 % | n/a | 0.0 % | 0.0 % | 0.0 % | 0.0 % | 0.0 % | 0.1 % | 0.0 % | 0.1 % | 1.1 % | n/a |
| W | 1.5 % | 0.0 % | 0.0 % | 0.2 % | 0.7 % | 0.8 % | 1.5 % | n/a | 3.2 % | 3.3 % | 0.0 % | 0.0 % | 1.6 % | 0.2 % | 0.0 % | 0.1 % | 1.0 % | n/a |
| X | 1.8 % | 0.7 % | 0.7 % | 2.0 % | 0.9 % | 1.0 % | 0.9 % | n/a | 0.0 % | 0.2 % | 0.0 % | 0.0 % | 0.0 % | 0.2 % | 0.0 % | 0.1 % | 1.7 % | n/a |
| other | 0.3 % | 2.7 % | 2.7 % | 5.7 % | 0.3 % | 1.4 % | 0.6 % | n/a | 0.2 % | 0.0 % | 0.0 % | 1.5 % | 0.8 % | 2.5 % | 0.3 % | 1.4 % | 1.2 % | n/a |

### Pop1 White: English/Welsh/Scottish/Northern Irish/British

These are frequencies from majority ethnic Britons, largely from rural areas, derived from over 4600 individuals (Helgason *et al.*, 2001, Røyrvik, unpublished).

### Pop2 White: Irish

Data from 300 individuals, from (Achilli *et al.*, 2007), originially from (McEvoy *et al.*, 2004).

### Pop3 White: Gypsy or Irish Traveller

Same as Pop3, given the lack of qualitative genetic differentiation between Traveller and non-Traveller Irish (Relethford and Crawford, 2013). Note that this category does not include Roma.

### Pop4 White: Other White

These frequencies are composed of 50% pan-European average frequencies and 50% Polish frequencies, to take account of the large Polish communities in London (see http://www.migrationobservatory.ox.ac.uk/briefings/migrants-uk-overview) and Birmingham, where Poles are 29% of this category. Variations in upper-hierarchy haplogroup frequencies are comparatively minor within Europe, and the New World white populations – derived as they primarily are from European parental populations – deviate little from European averages. Data from (Achilli *et al.*, 2007), comprising 10 970 individuals.

**(For ‘Mixed/multiple ethnic group’ categories, see below.)**

### Pop9 Asian/Asian British: Indian

This is a weighted composite of different frequencies from 2126 individuals from the Indian subcontinent (Dubut *et al.*, 2009), based on major regions from which immigrants to the UK originated (Chanda and Ghosh, 2013). Weighting was as follows: Punjab – 45%, Kerala, Tamil Nadu, Andra Pradesh and Gujarat – 13.75% each.

### Pop10 Asian/Asian British: Pakistani

This unweighted composite of 818 individuals includes all Pakistani groups from (Quintana-Murci *et al.*, 2004), as well as Indian Punjabis and Kashmiris from (Dubut *et al.*, 2009), as people hereditarily from the latter two geographic regions constitute by far the largest Pakistani sub-populations in the UK (TheChangeInstitute, 2009).

### Pop11 Asian/Asian British: Bangladeshi

From 30 individuals from (Dubut *et al.*, 2009).

### Pop12 Asian/Asian British: Chinese

These frequencies are from 2398 southern Chinese individuals from (Xue *et al.*, 2008), as the majority of immigration has traditionally been from southern China (Pharaoh, 2009). While this has perhaps changed (Pharaoh, 2009), for the purposes of our study, the differences in haplogroup composition, at our level of refinement, would not change materially.

### Pop13 Asian/Asian British: Other Asian

Defined as 50% Pop9, 50% Pop12. While the composition/ethnicity of different ‘other’ Asian groups is unknown, the Chinese and Indian samples, taken together, should for our purposes adequately represent the majority of East Asia, South Asia and Southeast Asia. Central Asia, with its higher frequencies of typically `West Eurasian’ haplogroups, may be underrepresented, but at a small fraction of 4.9-2.9%, this will not substantially affect our calculations, especially as these haplogroups are well represented in Pops1-4.

### Pop14 Black/African/Caribbean/Black British: African

This is a weighted composite of different frequencies from the African continent, scaled as 55% Western and Central African, 41% South and East African, and 3% North African (see (Owen, unpublished) for details on the origins of African immigrants to the UK). Sources for African regional and national haplogroup frequencies are (Fendt *et al.*, 2012, Kivisild *et al.*, 2004, Plaza *et al.*, 2003, Salas *et al.*, 2002, Saunier *et al.*, 2009). Note that North Africans are also included in pop17.

### Pop15 Black/African/Caribbean/Black British: Caribbean

From (Deason *et al.*, 2012), 400 individuals representing Jamaica. The vast majority of haplogroups are typical of Sub-Saharan Africa, but a small minority are likely the result of European (H, J, U) and Taino (A, B, D) ancestry.

### Pop16 Black/African/Caribbean/Black British: Other

These frequencies are composed of 50% pop14 and 50% pop15.

### Pop 17 Other ethnic group: Arab

Base frequencies from (Badro *et al.*, 2013) for 3248 individuals. Haplogroup frequencies are weighted averages of 33.3% each from the Arabian Peninsula, the Middle East, and Africa north of the Sahara.

### Pop18 Other ethnic group: Other

As there is no basis for estimating haplogroup frequencies for this category (2.1% in London, 1.0% in Birmingham), it has not been included in the analysis.

### Pop5 Mixed/multiple ethnic group: White and Black Caribbean

These frequencies are composed of 50% Pop1 and 50% Pop15. This assumes that the census respondents’ mothers are equally likely to be ‘White’ or ‘Black Caribbean’, which may not be the case.

### Pop6 Mixed/multiple ethnic group: White and Black African

These frequencies are composed of 50% Pop1 and 50% Pop14. This assumes that the census respondents’ mothers are equally likely to be ‘White’ or ‘Black African’, which may not be the case.

### Pop7 Mixed/multiple ethnic group: White and Asian

These frequencies are composed of 50% Pop1 and 50% Pop13. This assumes that the proportion of census respondents’ mothers are equally likely to be ‘White’ or ‘Asian’, which may not be the case.

### Pop8 Mixed/multiple ethnic group: Other Mixed

As there is no basis for estimating haplogroup frequencies for this category (1.4% in London, 0.8% in Birmingham), it has not been included in the analysis.

Note that minority haplogroups from any given sample population are often not assayed in the original source, so the ‘other’ category by necessity has some small overlap with the other haplogroups in the table for different groups. The maximum proportion of ‘other’ haplogroups is just 1.27% (in London), so it is not expected to bias the overall dataset unduly.

